# The long non-coding RNA *SAMMSON* is essential for uveal melanoma cell survival

**DOI:** 10.1101/2021.01.18.427176

**Authors:** Shanna Dewaele, Louis Delhaye, Boel De Paepe, Eric De Bony, Jilke De Wilde, Katrien Vanderheyden, Jasper Anckaert, Nurten Yigit, Eveline Vanden Eynde, Fariba Nemati, Didier Decaudin, Aart Jochemsen, Eleonora Leucci, Jo Vandesompele, Jo Van Dorpe, Chris Marine, Rudy Van Coster, Sven Eyckerman, Pieter Mestdagh

**Author notes:** Corresponding author Pieter Mestdagh, Corneel Heymanslaan 10, B-9000 Ghent, Belgium. Authors’ Contributions, P.M. conceived and supervised the project; S.D, L.D., B.D.P., E.D.B., J.D.W., K.V., N.Y, E.V.E. and S.E. designed and performed experiments; P.M., S.D., L.D., E.D.B., J.D.W., J.A analyzed the data; F.N., E.L. performed PDX experiments; A.J. provided cell lines; D.D., J.V.D., C.M, J.V., R.V.C., S.E. contributed technical support and resources; S.D. and P.M wrote the paper. All authors contributed to manuscript editing and approved the final draft. **Statement of translational relevance** Despite localized treatment, 50% of uveal melanoma (UM) patients develop metastasis. For these patients, currently no effective treatments are available, resulting in a median survival time of less than 1 year. We demonstrate that the melanoma specific lncRNA *SAMMSON* is consistently expressed in UM tumors and that ASO-mediated *SAMMSON* knockdown abrogates cell viability and induces apoptosis of UM cells derived from both primary and metastatic UM tumors, independent of their genetic background. Furthermore, subcutaneous injection of a *SAMMSON* ASO in different uveal melanoma PDX models significantly impairs in vivo tumor growth. Together, our findings indicate that *SAMMSON* is an attractive therapeutic target in UM and that ASO-mediated *SAMMSON* targeting in vivo elicits therapeutic effects in PDX models.

## Abstract

**Purpose:** Long non-coding RNAs (lncRNAs) can exhibit cell-type and cancer-type specific expression profiles, making them highly attractive as therapeutic targets. Pan-cancer RNA sequencing data revealed broad expression of the *SAMMSON* lncRNA in uveal melanoma (UM), the most common primary intraocular malignancy in adults. Currently, there are no effective treatments for UM patients with metastatic disease, resulting in a median survival time of 6-12 months. We aimed to investigate the therapeutic potential of *SAMMSON* inhibition in UM.

**Experimental Design:** The impact of antisense oligonucleotide (ASO)-mediated *SAMMSON* inhibition was evaluated in a panel of UM cell lines and patient derived xenograft (PDX) models. Cell proliferation and apoptosis were quantified *in vitro* and *in vivo* and complemented with molecular profiles established through RNA-sequencing. *SAMMSON* interaction partners were identified using ChIRP-MS.

**Results:** *SAMMSON* inhibition impaired the growth and viability of a genetically diverse panel of uveal melanoma cell lines. These effects were accompanied by an induction of apoptosis and were recapitulated in two uveal melanoma PDX models through subcutaneous ASO delivery. *SAMMSON* pulldown revealed several candidate interaction partners, including various proteins involved in mitochondrial translation. Consequently, inhibition of *SAMMSON* impaired global protein translation levels and mitochondrial function in uveal melanoma cells.

**Conclusion:** *SAMMSON* expression is essential for uveal melanoma cell survival. ASO-mediated silencing of *SAMMSON* may provide an effective treatment strategy to treat primary and metastatic uveal melanoma patients.

## Introduction

Uveal melanoma (UM) is the most common primary intraocular malignancy in adults, with an incidence of 5 - 7.4 cases per million annually(1–3). Current treatments consist of radiotherapy and enucleation, but despite the advances in local therapy towards eye-preserving therapeutic choices, no substantial progress in overall survival has been achieved. The main cause of death of UM patients is due to the metastatic dissemination, mainly to the liver, in about 50% of the patients. Patients with metastatic UM have extremely poor survival, with a median survival time of 6 - 12 months(4). Uveal melanoma is a genetically and biologically distinct type of melanoma that arises from choroidal melanocytes in the choroidal plexus, ciliary body and iris of the eye. In addition, UM and cutaneous melanoma differ in their chromosomal aberrations and mutational signature. Recurrent genomic aberrations in UM include loss of 1p, monosomy of chromosome 3, loss of 6q and 8p and gain of 6p and 8q(5,6). Tumor cells are characterized by activated G protein-coupled receptor (GPCR) signaling, which occurs in almost all UM tumors by specific mutations that have been related to UM, such as mutations in *GNAQ or GNA11* (90% of UM tumors). These mutations result in the activation of the pathways downstream of Gα_q_ and Gα_11_ such as the Ras Homolog Family Member/Ras-related C3 botulinum toxin/Yes Associated Protein (Rho/Rac/YAP) pathway, Phosphatidylinositol (4,5)-bisphosphate 3-kinase (PI3K)/AKT pathway and Phospholipase C (PLC) which subsequently activates the Mitogen-Activated Protein Kinase (MAPK) pathway (RAF/MEK/ERK)(7).

Long non-coding RNAs are an emerging class of regulatory RNA molecules that interact with proteins, DNA, and other RNA molecules. Compared to protein-coding mRNAs, lncRNAs show a more tissue restricted expression profile, making them attractive targets for therapy. LncRNAs are regulating a variety of cellular functions such as transcription, splicing, mRNA stability and translation, and do so via different mechanisms. Recently, Survival Associated Mitochondrial Melanoma Specific Oncogenic Non-coding RNA (*SAMMSON*) was discovered on chromosome 3p13 as a lineage survival oncogene in skin melanoma(8). Interaction of *SAMMSON* with proteins involved in ribosomal RNA (rRNA) maturation and protein synthesis such as p32 (C1QBP), CARF and XRN2(8,9) and sequestration of CARF in the cytoplasm result in an elevated cytosolic and mitochondrial rRNA processing and protein synthesis(9). In this study, we show that *SAMMSON* is consistently expressed in UM and conjunctival melanoma (CM) cells, the latter of which are phenotypically and genetically more related to skin melanoma. *SAMMSON* silencing, by means of locked nucleid acid (LNA) antisense oligonucleotides (ASOs), revealed an essential role for *SAMMSON* in UM survival *in vitro* and *in vivo* and provides perspectives for RNA-targeted therapy.

## Material and methods

### Cell culture

For all human cell lines, an ethical approval was obtained from the Ghent University commission for medical ethics. The human uveal melanoma cell lines 92.1(10), OMM1(11), OMM2.3, MEL270, MP38(12), MP46(12), MM28(12) and MP65(12) and conjunctival melanoma cell lines CRMM1 and CRMM2(13) were obtained from the Leiden University Medical Center, The Netherlands. OMM2.3 and MEL270 were a kind gift from Bruce Ksander. MM28, MP65, MP46 and MP38 were a kind gift from Sergio Roman-Roman. Uveal melanoma cell line MEL077 and skin melanoma cell line SK-MEL28 was obtained from the Laboratory for Molecular Cancer Biology, VIB-KU Leuven, Belgium. *SAMMSON* negative cell lines HEK293T and CT5.3hTERT were obtained from ATCC and the Department of Radiation oncology and experimental cancer research, Ghent University, Belgium, respectively. The cell lines 92.1, OMM1, OMM2.3 and MEL270 were grown in Dulbecco’s modified Eagle’s medium (DMEM, Gibco)/F12 – Roswell Park Memorial Institute (RPMI, Gibco) 1640 (1:1) medium. SK-MEL28 was grown in DMEM/F-12 GlutaMAX medium (Gibco), CT5.3hTERT in DMEM medium, CRMM1 and CRMM2 in Ham’s F-12K (Kaighn’s) medium (Gibco), MM28, MP65, MP46 and MP38 in Iscove’s Modified Dulbecco’s Medium (IMDM) and HEK293T and MEL077 in RPMI 1640 medium. Media of MM28, MP65, MP46 and MP38 were supplemented with 20% fetal bovine serum (FBS) and all other media with 10% FBS, 2 mM L-glutamine (Gibco) and 100 IU/ml penicillin/streptomycin (Gibco) and all cell lines were incubated in a humidified atmosphere containing 5% CO_2_ at 37°C. For executing experiments, cells were grown in L-glutamine and penicillin/streptomycin free media. Short tandem repeat (STR) genotyping was used to validate cell line authenticity and absence of mycoplasma was verified on a monthly basis for all cell lines in culture.

### Animal models

The *in vivo* experiment with PDX model MEL077(14) was executed at the Patient Derived Tumor Xenograft Platform, Trace, Leuven, Belgium. The experimental protocol and animal housing were in accordance with institutional guidelines as put forth by the Belgian Ethical Committee and the KU Leuven animal ethics committee that approved this study (agreement P164/2019, KU Leuven, Belgium). Female Naval Medical Research Institute (NMRI) mice were engrafted at an age of 12 weeks with an UM tumor fragment of 3 mm^3^. Mice bearing tumors with a volume of 150 mm^3^ were individually identified and randomly assigned to the different treatment/control groups. Mice were weighed twice a week. Tumor volumes were calculated by measuring every 2 days the length (L), width (W) and height (H) of the tumor with calipers. Each tumor volume (V) was calculated according to the following formula: V = (LxWxH)x(π/6). At the end of treatment, mice were sacrificed with collection of tumor, liver, lung and blood. ASO 3 and NTC ASO were injected subcutaneously at a dose of 10 mg/kg 3.5 times a week. An independent repeat of this experiment has been performed using an independent mouse cohort. The *in vivo* experiment with PDX model MP46 (BAP1 negative, GNAQ^mut^, monosomy 3) was executed at the Institut Curie, Paris, France. Animal care and use for this study were performed in accordance with the recommendations of the European Community (2010/63/UE) for the care and use of laboratory animals. Experimental procedures were specifically approved by the ethics committee of the Institut Curie CEEA-IC #118 (Authorization APAFiS# 25870-2020060410487032-v1 given by National Authority) in compliance with the international guidelines. Female severe combined immunodeficiency (SCID) mice were engrafted with an UM tumor fragment of 15 mm^3^. Mice bearing tumors with a volume of 60 - 180 mm^3^ were individually identified and randomly assigned to the control or treatment groups. Mice were weighed and tumors were measured twice a week. Tumor volumes were calculated by measuring two perpendicular diameters with calipers. Xenografted mice were sacrificed at the end of the treatment. Each tumor volume (V) was calculated according to the following formula: V = a × b^2^/2, where a and b are the largest and smallest perpendicular tumor diameters. ASO 3 and NTC ASO were administered subcutaneously at a dose of 10 mg/kg 3 times a week in the first week of treatment and twice a week in the following weeks.

Relative tumor volumes (RTV) were calculated with the following formula: RTV = (Vx/V1), where Vx and V1 are the tumor volumes on day x and the first day of treatment, respectively.

### Compounds and antisense oligonucleotides (ASOs)

Tigecycline was purchased from Selleckchem.

The LNA GapmeR oligonucleotides specifically targeting *SAMMSON* and the LNA non-targeting control (NTC) GapmeR were purchased from Qiagen (catalog number 339517).

Sequence LNA ASO 3: GTGTGAACTTGGCT

Sequence LNA ASO 11: TTTGAGAGTTGGAGGA

Sequence LNA NTC: TCATACTATATGACAG

### Cell proliferation and apoptosis assay

Cells were seeded in 96 well plates (Corning costar 3596) at a density of 5000 cells/well and were allowed to settle overnight. Subsequently, the cells were transfected with ASO 3 or NTC ASO using lipofectamine 2000 (Thermo Fisher Scientific) or TransIT-X2 (Mirus Bio).

Cell viability and apoptosis were examined using a CellTiter-Glo assay (Promega) and Caspase 3/7 assay (Promega), respectively. Before initiating the assay, the culture plates and reconstituted assay buffer were placed at room temperature for 30 minutes. Next, the culture medium was replaced by 200 μl fresh culture medium - assay buffer (1:1) mix. To induce complete cell lysis, the plates were shaken during 10 min. 100 μl from each well was subsequently transferred to an opaque 96-well plate (Nunc), which was measured with a GloMax 96 Microplate Luminometer (Promega).

For real-time analysis, the IncuCyte Zoom system and IncuCyte S3 system (Essen BioScience) were used. Cells were seeded and treated as described above. After treatment, the culture plate was incubated in an IncuCyte Zoom system or IncuCyte S3 system at 37°C in a humidified 5% CO2 incubator. Phase contrast whole well images were captured every 3 h. The IncuCyte ZOOM (version 2016B) and IncuCyte S3 software (version 2019B) (Essen BioScience) were utilized in real-time to measure % confluence, as a proxy for proliferation. The % confluence values were corrected for seeding variability resulting in normalized % confluence values. For apoptosis assessment, annexin V (Essen BioScience, dilution 1:400) was added while treating the cells. The IncuCyte ZOOM and IncuCyte S3 software (Essen BioScience) were utilized in real-time to measure % confluence and total Red object area (μm^2^/well). The ratio between total Red object area and phase object confluence (%) is resulting in relative annexin V ratios.

### SUnSET

Cells were seeded in T75 culture flasks (Cellstar) at a density of 1.17 × 10^6^ cells/flask 24h prior to transfection. The cells were transfected with 50 nM ASO 3 or NTC ASO using lipofectamine 2000. After 24h, cells were washed in 1x phosphate buffer saline (PBS) and subsequently incubated with puromycin containing media (InvivoGen, 10 μg/ml) for 10 min. Puromycin incorporation is a proxy for the mRNA translation rate *in vitro* and was measured by western blotting using an anti-puromycin antibody (MABE343, clone 12D10, Merck Millipore, 1:10 000). The antibody was diluted in Milk/TBST (5% non-fat dry milk in TBS with 0.1% Tween20). A ponceau S staining (Sigma Aldrich) was performed to verify equal loading.

### Western blot analysis

Cells were lysed in RIPA lysis buffer (5 mg/ml sodium deoxycholate, 150 mM NaCl, 50 mM Tris-HCl pH 7.5, 0,1% SDS solution, 1% NP-40) supplemented with protease and/or phosphatase inhibitors. Protein concentrations were determined with the BCA protein assay (Bio-Rad). In total, 35 μg of protein lysate was loaded onto an SDS-PAGE gel (10% Pre-cast, Bio-Rad), ran for 1 h at 100 V and subsequently blotted onto a nitrocellulose membrane. HRP-labeled anti-rabbit (7074 S, Cell Signaling, 1:10000 dilution) and anti-mouse (7076P2, Cell Signaling, 1:10000 dilution) antibodies were used as secondary antibodies. The antibodies were diluted in Milk/TBST (5% non-fat dry milk in TBS with 0,1% Tween20) and antibody binding was evaluated using the SuperSignal West Dura Extended Duration Substrate (ThermoFisher Scientific) or SuperSignal West Femto Maximum Sensitivity Substrate (ThermoFisher Scientific). Imaging was done using the Amersham Imager 680 (GE Healthcare). Image J (version 1.52q) was used for the quantification of the blots. Uncropped scans of the blots can be found in Supplemental Fig 3C.

### Reverse transcription quantitative polymerase chain reaction (RT-qPCR)

Total RNA was extracted using the miRNeasy kit (Qiagen) according to the manufacturer’s instructions, including on-column DNase treatment. The Nanodrop (ThermoFisher Scientific) was used to determine RNA concentrations and cDNA synthesis was performed using the iScript Advanced cDNA synthesis kit (Bio-Rad) using a mix containing 200 ng of RNA, 4 μl of 5x iScript advanced reaction buffer and 1 μl of iScript advanced reverse transcriptase. The qPCR reactions contain 2 μl of 1:4 diluted cDNA (2.5 ng/μl), 2.5 μl SsoAdvanced Universal SYBR Green Supermix (Bio-Rad), 0.25 μl forward (5 μM, IDT) and 0.25 μl reverse primer (5 μM, IDT) and was analyzed on a LC480 instrument (Roche).

For some purposes RNA was obtained using the SingleShot Cell Lysis Kit (Bio-Rad) according to the manufacturer’s instructions and cDNA synthesis was performed using the iScript Advanced cDNA synthesis kit (Bio-Rad) using a mix containing 8 μl sample lysate, 7 μl nuclease free water, 4 μl of 5x iScript advanced reaction buffer and 1 μl of iScript advanced reverse transcriptase. Subsequently, the cDNA is diluted 4 times prior to the qPCR reaction (see higher).

Expression levels were normalized using expression data of at least 2 stable reference genes out of 4 tested candidate reference genes (SDHA, HPRT1, UBC and TBP). Multi-gene normalization and relative quantification was performed using the qbase+ software (v3.2, www.qbaseplus.com).

The primer sequences used for qPCR were as follows:

*SAMMSON* Fw: CCTCTAGATGTGTAAGGGTAGT, Rv: TTGAGTTGCATAGTTGAGGAA

SDHA Fw: TGGGAACAAGAGGGCATCTG, Rv: CCACCACTGCATCAAATTCATG

HPRT1 Fw: TGACACTGGCAAAACAATGCA, Rv: GGTCCTTTTCACCAGCAAGCT

UBC Fw: ATTTGGGTCGCGGTTCTTG, Rv: TGCCTTGACATTCTCGATGGT

TBP Fw: CACGAACCACGGCACTGATT, Rv: TTTTCTTGCTGCCAGTCTGGAC

### Quantitative PCR for evaluation of metastatic disease

For assessment of metastatic disease in UM PDX models MEL077 and MP46, whole blood, lung and liver tissues were collected at the end of the experiment of ASO 3 or NTC ASO treated mice. Genomic DNA was isolated from liver and lung tissues (n=4 for MEL077 and n=6-7 for MP46 per treatment group) using the QIAamp DNA Mini Kit (Qiagen) and for blood (n=3-4 for MEL077 and n=6-7 for MP46 per treatment group) using the QIAamp DNA Blood Mini kit (Qiagen), according to the manufacturer’s instructions. The DropSense 96 (Trinean) was used to determine DNA concentrations. The qPCR reactions contain 2 μl of gDNA (6 ng/μl), 2.5 μl SsoAdvanced Universal SYBR Green Supermix (Bio-Rad), 0.25 μl forward (5 μM, IDT) and 0.25 μl reverse primer (5 μM, IDT) and was analysed on a LC480 instrument (Roche).

Copy number levels were determined for the human Alu-Sq, SVA and LINE-1 repetitive DNA sequences and murine gDNA assays located in the Hprt1 and Pthlh genes were used as reference genes for normalization. Analysis was performed using the qbase+ software (v3.2, www.qbaseplus.com). qPCR results from both PDX experiments were combined after log transformation, mean centering and autoscaling (according to Willems et al. (15)). For all human repetitive DNA sequence assays, high Cq values were obtained in the negative qPCR control. Five Cq values difference between the samples from the NTC ASO treated mice and the negative qPCR control was taken as a cutoff to conclude the presence of human DNA. Only lung samples fulfilled the criteria.

The oligonucleotide primers used for qPCR were as follows:

Alu-Sq Fw: CATGGTGAAACCCCGTCTCTA, Rv: GCCTCAGCCTCCCGAGTAG

SVA Fw: CTGTGTCCACTCAGGGTTAAAT, Rv: GAGGGAAGGTCAGCAGATAAAC

LINE-1 Fw: TGGCACATATACACCATGGAA, Rv: TGAGAATGATGGTTTCCAATTTC

Hprt1 Fw: CCTAAGATGAGCGCAAGTTGAA, Rv: CCACAGGACTAGAACACCTGCTAA

Pthlh: GACGTACAAAGAACAGCCACTCA, Rv: TTTTTCTCCTGTTCTCTGCGTTT

### RNA immunoprecipitation (RIP)

20 × 10^6^ 92.1 cells were harvested from T75 culture flasks (Cellstar) using trypsin, followed by a 5 min. centrifugation step at 500 g at 4 °C. After removal of the supernatant, cell pellets were resuspended in 1 ml cold PBS in 1.5 ml tube, followed by a second 5 min. centrifugation step at 600 g at 4 °C. After supernatant removal, pellets were flash frozen and stored at −80 °C until use. Pellets were lysed in 4 ml of polysome buffer (for 100 ml: 2 ml of TRIS 1 M pH 8.0, 4 ml of NaCl 5 M, 250 μl of MgCl2 50 μl of Triton 1 M and 100 μl of DTT 1 mM. Add fresh, 250 μl of RNAsin, (Promega), 1 ml of vanadyl ribonucleoside complexe solution and 2.5 ml of protease inhibitors, (Sigma)) for 30 minutes on ice. Lysates were pre-cleaned with 20 μl of dynabeads (Life Technologies) per sample for 1 h at 4 °C with rotation. No antibody, 5 μg of human IgG (Abcam, ab2410) or 5 μg of XRN2 (Bethyl laboratories, A301-103A) and p32 antibodies (Bethyl Laboratories, A302-863A) were added to lysates and incubated overnight at 4 °C. Antibody-protein complexes were pulled down with 100 μl of rinsed beads per sample during 1 hour at 4 °C for 4 hours with rotation. Beads were captured on a magnetic rack and rinsed 5 times with polysome buffer. Rinsed beads were then resuspended in Qiazol (Qiagen) for RNA extraction following the manufacturer’s instructions. Finally, 1 μl of sequin spike (https://www.sequinstandards.com) was added to 14 μl of RNA for cDNA synthesis with SsoAdvanced iScript (Bio-Rad), followed by RT-qPCR using SsoAdvanced master mix (Bio-Rad). *SAMMSON* Cq values were normalized to sequin Cq values using the qbase+ software (v3.2, www.qbaseplus.com).

The primer sequences used were as follows:

*SAMMSON* Fw: CCTCTAGATGTGTAAGGGTAGT, Rv: TTGAGTTGCATAGTTGAGGAA

Sequin Fw: ATGCTTTGATCGCGTTGGTG, Rv: AGCAAAACGAACGGACAATGA

### ChIRP-MS affinity purification

75 × 10^6^ – 100 × 10^6^ cells were cultured in 145 cm^2^ dishes at a maximum confluency of 80%, washed once with ice-cold PBS, and UV cross-linked in ice-cold PBS at 254 nm to an accumulated energy intensity of 400 mJ/cm^2^. Cells were scraped in ice-cold PBS, and split equally among eight microcentrifuge tubes. ChIRP lysis buffer (20 mM Tris-HCl pH 7.5, 200 mM NaCl, 2.5 mM MgCl_2_, 0.05% NP-40, 0.1% SDS) (8) was supplemented with fresh 0.1% sodiumdeoxycholate, 60 U/mL Superase-In Rnase inhibitor (Invitrogen), 1 mM DTT, 0.5 mM PMSF, and protease inhibitor cocktail (Roche). Cell pellets were resuspended in supplemented ChIRP lysis buffer, and sonicated with a Bioruptor (Diagenode) until lysates appeared clear. 10% of the ChIRP sample was used for RNA extraction of input material. Thereafter, 6.23 μl of 50 μM *SAMMSON* or LacZ biotinylated capture probes (LGC Biosearch Technologies) were bound to 100 μl of equilibrated Rnase-free Dyna-One C1 magnetic beads (Thermo) per sample and were incubated overnight at 4 °C with end-to-end rotation. Next day, *SAMMSON* or LacZ probe-bound beads were added to the lysates and lysates were rotated for 3 h at 4 °C. Bead-bound fractions were washed three times with unsupplemented ChIRP lysis buffer. 10% of the sample was used for RNA extraction to validate RNA pulldown on RT-qPCR. Next, beads were washed three times with Rnase-free trypsin digestion buffer (20 mM Tris-HCl pH 7.5, 2 mM CaCl_2_), and were ultimately resuspended in 20 μl 20 mM Tris-HCl pH 7.5. 750 ng trypsin was added directly on the beads, and digestion was left overnight at 37 °C. Next day, an additional 250 ng trypsin was added and incubated for 3 h at 37 °C. Peptides were acidified to a final concentration of 2% formic acid. All experiments were performed in biological triplicates for label-free quantitative proteomic analysis.

### LC-MS/MS instrument analysis

Peptide mixtures were run on an Ultimate 3000 RSLC nano LC (Thermo Fisher Scientific) connected in-line to a Q-Exactive HF mass spectrometer (Thermo Fisher Scientific). In brief, peptides were loaded on an in-house made trapping column (100 μm i.d. × 20 mm, 5 μM C18 Reprosil-HD beads, Dr. Maisch, Ammerbuch-Entringen, Germany). After flushing the trapping column, peptides were loaded in solvent A (0.1% formic acid) on an in-house made reverse-phase column (75 μm i.d. × 250 mm, 3 μm Reprosil-Pur-basic-C18-HD beads packed in the needle, Dr. Maisch, Ammerbuch-Entringen, Germany) and eluted by a linear gradient of solvent B (0.1% formic acid in acetonitrile) from 2% to 55% in 1.5 h, and subsequently washed with 99% solvent B. All steps were run at a constant flow rate of 300 nl/min. The mass spectrometer was operated in a data-dependent acquisition, positive ionization mode, automatically switching between MS and MS/MS acquisition for the five most abundant peaks in a MS spectrum. Source voltage was 3.4 kV, and capillary temperature was 275 °C. One MS1 scan (m/z 400-2000, AGC target 3 × 106 ions, maximum ion injection time 80 ms), acquired at a resolution of 70 000 (at 200 m/z), was followed by up to five tandem MS scans (resolution 17 500 at 200 m/z) of the most intense ions, fulfilling the predefined selection criteria (AGC target 5 × 104 ions, maximum ion injection time 80 ms, isolation window 2 Da, fixed first mass 140 m/z, spectrum data type: centroid underfill ratio 2%, intensity threshold 1.3 × 104, exclusion of unassigned 1, 5 - 8, and >8 positively charged precursors, peptide match preferred, exclude isotopes on, dynamic exclusion time 12 ms). The HCD collision energy was set to 25% normalized collision energy, and the poly(dimethylcyclosiloxane) background ion at 445.120025 Da was used for internal calibration (lock mass).

### MaxQuant and Perseus MS data processing and analysis

Xcalibur raw files were analysed using the Andromeda search engine implemented in MaxQuant (MaxQuant v1.6.0.1). Spectra were searched against the human UniProt sequence database. Methionine oxidation and N-terminal acetylation were set as variable modifications. The minimum label-free quantitation ratio count was 2, and the Fast LFQ option was disabled. After the searches were completed, LFQ intensities were imported in Perseus (v1.5.8.5) for downstream analysis.

LFQ intensities were log 2 transformed, and contaminant proteins, reverse hits, and protein only identified by site were excluded from the analysis. Three valid values in at least one sample group (i.e. pulldown of *SAMMSON* or LacZ) was used for a protein to be included in further analysis. Missing values were imputed from a normal distribution of intensities. A two-sided t-test (0.05 FDR, 1000 randomizations) was performed to identify differential proteins in volcano plots. The default S0-value (0.1) in Perseus was maintained for generating the S-curve in the volcano plots. The mass spectrometry proteomics data have been deposited to the ProteomeXchange Consortium via the PRIDE(16) partner repository with the dataset identifier PXD023511. Reactome (https//reactome.org/) was used to perform overrepresentation analysis.

### JC-1 fluorescent staining

Cells were seeded in Nunc Lab-Tek Chamber slides at a density of 100 000 cells/well 24 h prior to transfection. Cells were transfected with 100 nM ASO 3 or NTC ASO using lipofectamine 2000. Cells were stained with 5 μg/ml of 5,5′,6,6′-tetraethylbenzimidazolyl-carbocyanine iodide (JC-1; Invitrogen) for 30 min at 37 °C following a published procedure(17). Live cells were visualized under a fluorescence microscope (Olympus, Hamburg, Germany), detecting red and green fluorescent emission separately using optical filters. Red over green JC-1 fluorescence ratios were determined by converting images to bright field and measuring the average grayscale (Cell F software; Olympus) in ten microscopic fields ×400 magnification selected at random, and reported as mean values ± SD.

### Seahorse XF Cell Mito Stress Test

A seahorse XF Cell Mito Stress Test was performed to measure the oxygen consumption rate (OCR). Cells were seeded in T25 culture flasks (Cellstar) at a density of 390 000 cells 24 h prior to transfection. The cells were transfected with 100 nM ASO 3 or NTC ASO using lipofectamine 2000. Four hours later, 15 000 cells were transferred to Seahorse XFp Cell Culture Miniplates (Agilent) and were allowed to settle overnight. Subsequently, oxygen consumption rates were measured in triplicates for each condition using the Seahorse XFp device (Agilent) according to the standard mito stress test procedures in seahorse assay medium supplemented with 14.3 mM glucose, 1 mM pyruvate and 2 mM glutamine (Sigma), and cells were sequentially challenged with 1 μM oligomycin, 0.5 μM (92.1) or 1 μM (OMM2.3) carbonyl cyanide 4-(trifluoromethoxy)-phenylhydrazone (FCCP) and 0.5 μM of a rotenone antimycin A mix (Agilent). Following the assay, protein concentrations were calculated based upon absorbance reading at 280 nm (Biodrop, Isogen Lifescience) for normalization of the results. Spare respiratory capacity was calculated as the difference between maximal and the basal oxygen consumption rate. Relative oxygen consumption rates are relative compared to NTC ASO values.

### Differential gene expression analysis by RNA sequencing

RNA sequencing was performed on quadruplicates of NTC ASO or ASO 3 (50 nM) treated 92.1 and OMM1 cells. Libraries for RNA sequencing were prepared using the Quant Seq 3’end library prep according to the manufacturer’s instructions (Lexogen) (2.5 μl input of RNA lysates) and quantified on a Qubit Fluorometer prior to single-end sequencing with 75 bp read length on a NextSeq 500 sequencer (Illumina).

RNA sequencing was performed on UM PDX MEL-077 tumor samples treated with either NTC ASO or ASO 3 (10 mg/kg). Libraries for RNA sequencing were prepared using the Truseq mRNA library prep according to the manufacturer’s instructions (Illumina) (500 ng input of purified RNA) and quantified on a Qubit Fluorometer prior to single-end sequencing with 75 bp read length on a NextSeq 500 sequencer (Illumina). Reads were mapped to the human genome (hg38) using STAR and gene expression was quantified using HTSeq (v0.6.1). Differentially expressed genes were identified using DESeq2 (v1.26.0). Pre-ranked gene set enrichment analysis (GSEA 4.1.0) was performed using c2.all.v7.2.symbols (curated gene sets), h.all.v7.2.symbols (Hallmark gene sets) and c6.all.v7.2.symbols (oncogenic signature gene sets) (Molecular Signatures Database (MsigDB)), applying 1000 permutations and a classic enrichment statistic.

### Immunohistochemistry (IHC)

Tumor samples of UM PDX mice (MEL077) that have been treated with ASO 3 or NTC ASO (10 mg/kg, 3.5x per week) for 3 weeks were fixed in 4% formaldehyde and subsequently embedded in paraffin to obtain formalin-fixed, paraffin-embedded (FFPE) samples. Tumor sections were used for hematoxilin and eosin (HE) staining, and for immunohistochemistry with antibodies against Ki-67 (RTU, clone 30-9, Roche) and caspase-3 (1:2000, AF835, R&D Systems) on a Ventana Benchmark Ultra automated staining system (Roche). The presence of apoptotic bodies was semi-quantitatively scored by two independent pathologists, blinded to treatment allocation, as follows: not present (−), rare (+), easily recognized (++), abundant (+++). Based on these scores, a diagnosis was made as ‘ASO 3-treated’ or ‘NTC ASO-treated’ for each tumor sample. Ki-67 expression was assessed by identifying areas with the most intense nuclear staining (‘hot spots’) at low magnification (100x). The percentage of immunoreactive tumor cells was calculated by counting at least 500 cells within each hot spot. Additionally, the mitotic rate was determined by counting the number of mitotic figures in 10 random high-power fields (400x) on HE sections.

### Statistical analysis

Statistical analyses and data visualizations were performed with Graphpad Prism version 9.0.0 (GraphPad Software, San Diego, California USA).

## Results

### SAMMSON is consistently expressed in uveal melanoma tumors

Pan cancer RNA sequencing data from more than 10 000 tumor samples representing 32 cancer types (The Cancer Genome Atlas, TCGA) showed the highest and most consistent *SAMMSON* expression in skin melanoma (SKCM, *SAMMSON* expression in >90% of tumor samples) followed by uveal melanoma (*SAMMSON* expression in >80% of tumor samples) (Fig 1A). In uveal melanoma tumors, *SAMMSON* expression is independent from patient survival, tumor stage, tumor localization site (choroid, ciliary body or iris) and metastatic state (Supplemental Fig 1). *SAMMSON* expression was further verified by RT-qPCR in various UM cell lines originating from primary tumors (92.1 and MEL270) and metastatic tumors (OMM2.3 and OMM1) (Fig. 1B), as well as in UM PDX-derived cell lines (MP38 (BAP1 negative, monosomy 3), MP46 (BAP1 negative, monosomy 3), MEL077, MP65 (BAP1 negative, monosomy 3), and MM28)(Fig 1C). *SAMMSON* expression was also detected in the conjunctival melanoma (CM) cell lines CRMM1 and CRMM2 that are genetically and phenotypically more related to skin melanoma(18) (Fig 1B). In contrast to skin melanoma tumors, where *SAMMSON* is invariably co-amplified with Melanocyte Inducing Transcription Factor (MITF) on chromosome 3, uveal melanoma cells are characterized by frequent (50 - 60%) loss of an entire copy of chromosome 3 (6,12) (Figure 1E). While the majority of genes expressed from chromosome 3 (n=581, 89%, Supplemental Fig 2) are significantly downregulated in monosomy 3 UM tumors, *SAMMSON* expression is independent of chromosome 3 copy number (Fig 1D, p=0.20, Mann-Whitney test). This suggests a compensation mechanism by which UM tumors cells maintain high *SAMMSON* levels in the presence of chromosome 3 loss.

**Fig 1.**
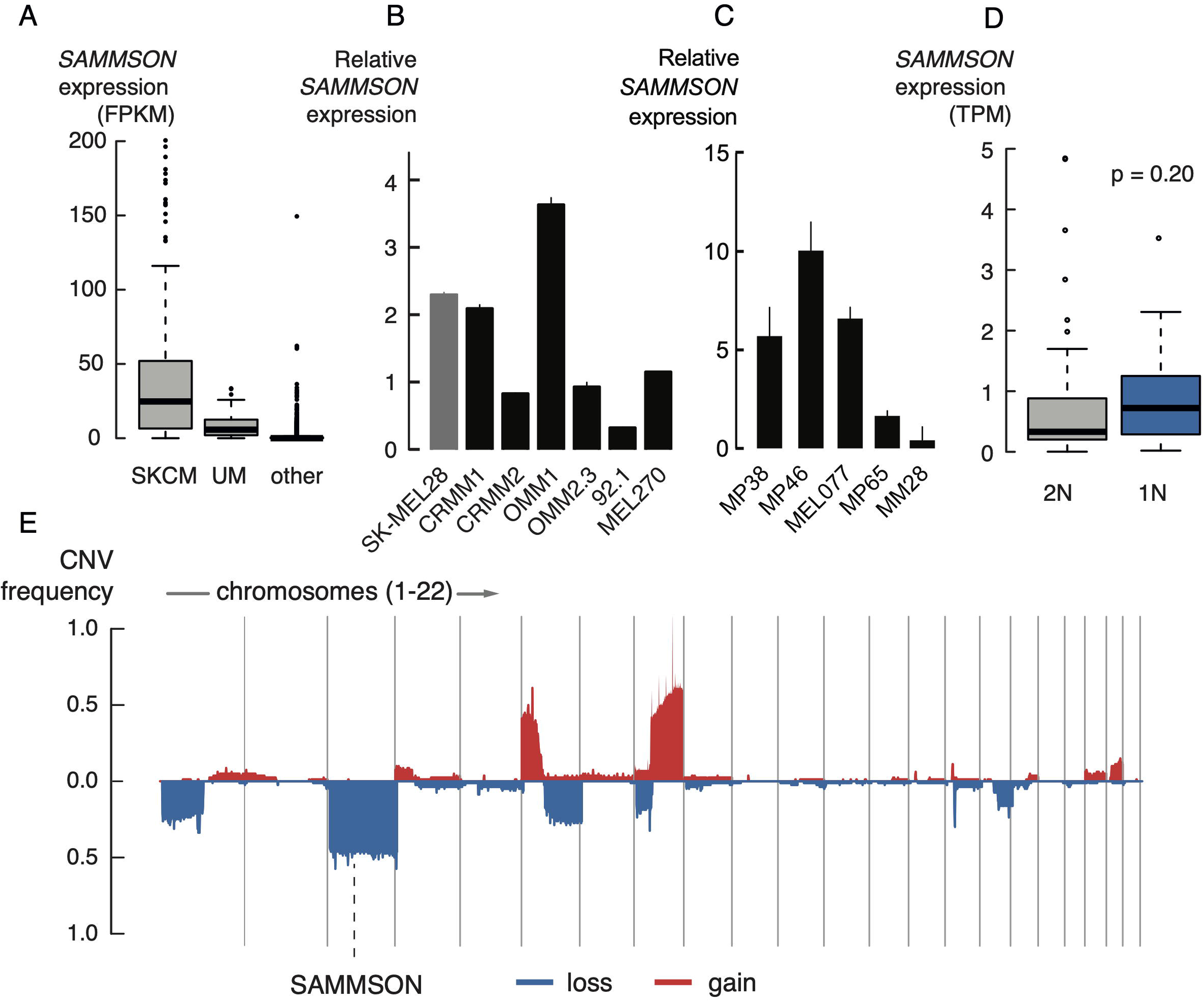
LncRNA *SAMMSON* is consistently expressed in UM tumors. **A.** RNA sequencing data from >10 000 tumor samples and 32 cancer types (TCGA) showing *SAMMSON* expression in skin melanoma (SKCM), uveal melanoma (UM) and other cancer types. **B.** Relative *SAMMSON* expression in CM cell lines (CRMM1, CRMM2) and UM cell lines (OMM1, OMM2.3, 92.1 and MEL270) compared to skin melanoma cell line SK-MEL28. Error bars represent ± standard error (SE) of qPCR replicates. **C.** Relative *SAMMSON* expression in UM PDX lines. Error bars represent ± standard error (SE) of qPCR replicates. **D.** No difference in *SAMMSON* expression levels between disomy 3 (2N, n=44) and monosomy 3 (1N, n=36) tumors (p=0.20, Mann-Whitney test). **E.** Frequent chromosomal aberrations in UM tumors and the location of *SAMMSON* on chromosome 3, which is frequently lost.

### SAMMSON expression is required for UM cell survival in vitro

To evaluate the importance of *SAMMSON* in UM and CM, we studied the effects of *SAMMSON* knockdown in various UM and CM cell lines using two independent *SAMMSON* targeting ASOs (ASO 3 and ASO 11). Transfection of 100 nM of both *SAMMSON* inhibiting ASOs significantly decreased *SAMMSON* expression with 60% and 66% in 92.1 and 37% and 36% in OMM2.3, respectively (Figure 2A, p<0.0001, one-way ANOVA). *SAMMSON* knock-down resulted in a strong and significant decrease in cell viability (based on ATP measurements) in 92.1 and OMM2.3 cells, as well as four other UM and two CM cell lines (Figure 2C, p≤0.001 for all except ASO 11 in MEL270, one-way ANOVA). These effects were accompanied by the induction of apoptosis, as evidenced by a significant increase in caspase-3/7 levels, observed for both *SAMMSON* inhibiting ASOs in 8 different UM and CM cell lines (Figure 2C, p≤0.05, one-way ANOVA). In line with earlier observations in skin melanoma cells(8), these effects were independent of the mutational status (GNAQ, GNA11, BAP1, BRAF and NRAS) of the UM and CM cell lines.

**Fig 2.**
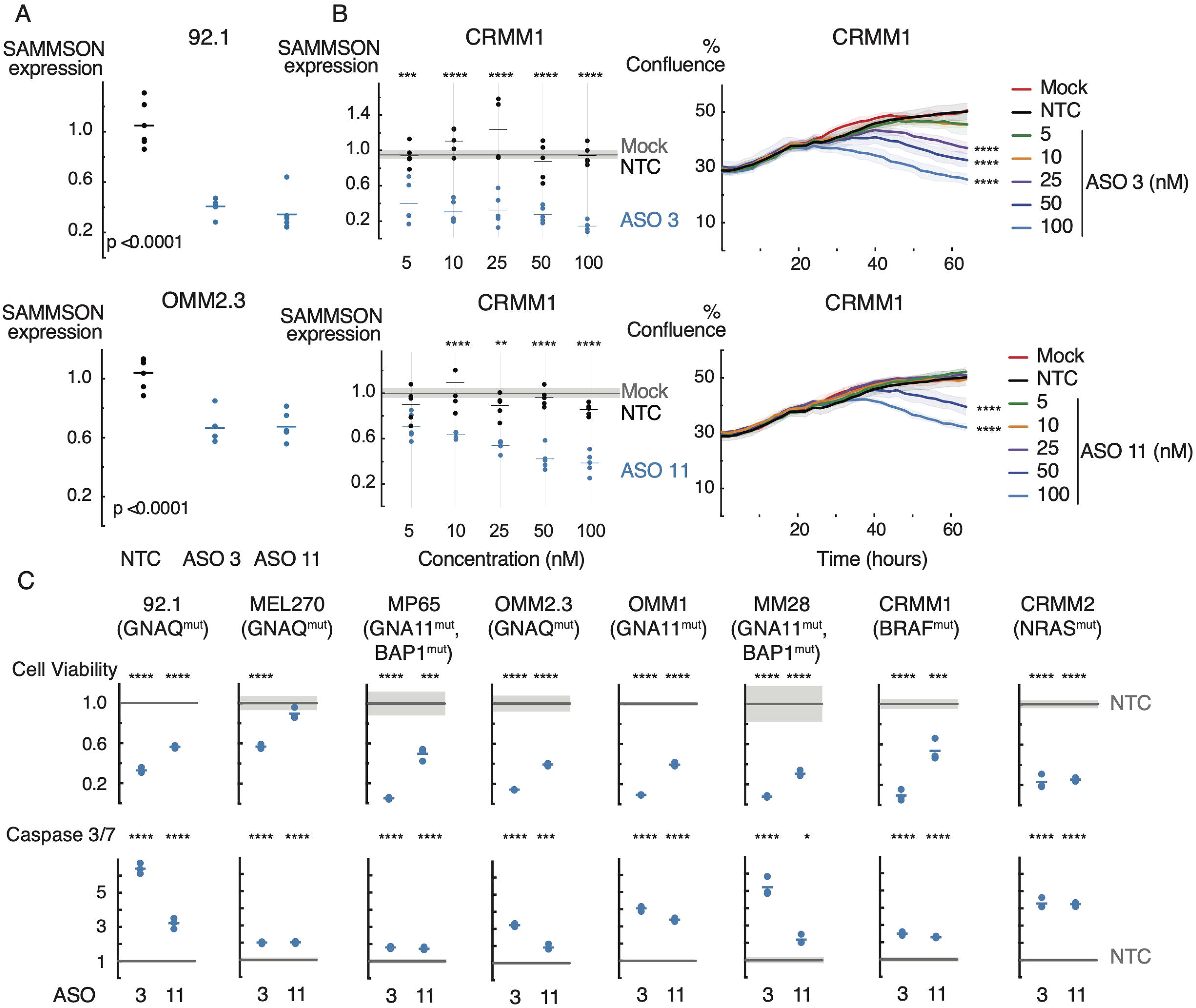
*SAMMSON* knockdown reduces cell viability and induces apoptosis. **A.** Relative *SAMMSON* expression in two UM cell lines transfected with 100 nM of a scrambled ASO (NTC), ASO 3 or ASO 11. The results are obtained 48h after transfection and represent the average of six replicates. **B.** Relative *SAMMSON* expression of CM cell line CRMM1 24h after transfection with 5, 10, 25, 50 or 100 nM of NTC ASO, ASO 3 or ASO 11 and scaled to untreated cells (Mock). Data-points represent 5 replicates. Reduction on proliferation in CRMM1 upon ASO 3 or ASO 11 treatment compared to NTC ASO treated and untreated cells (Mock) measured with time-lapse microscopy (every 2-3 h) using the IncuCyte device. Error bars represent ± s.d. of 3 replicates. **C.** Viability (Cell titer glo) and Caspase (Caspase-Glo 3/7) assays in NTC ASO, ASO 3 and ASO 11 treated UM and CM (CRMM1 and CRMM2) cells (72 h post-transfection). The data are presented as the mean of 3 replicates ± s.d. P-values were calculated using one-way ANOVA with Dunnett’s multiple testing correction.

To evaluate whether *SAMMSON* knockdown depends on ASO dosing, we performed a dose response experiment with both ASOs in the CM cell line CRMM1. We observed a dose dependent decrease in *SAMMSON* expression for both ASOs (Figure 2B, p≤0.001 for all tested concentrations of ASO 3 and p≤0.01 for all tested concentrations of ASO 11, except 5 nM, one-way ANOVA). We also observed a dose dependent reduction in cell growth and proliferation, measured via real-time cell imaging (Figure 2B, p≤0.0001 for the 3 and 2 highest concentrations using ASO 3 or ASO 11, respectively, one-way ANOVA).

To verify that the observed effects were not related to the lipid-based delivery of the ASOs, we evaluated an alternative non-lipid-based delivery method using the TransIT-X2 transfection reagent. Transfection of 100 nM of ASO 3 significantly decreased *SAMMSON* expression in 92.1 and OMM1 (Supplemental Fig 3A, p<0.0001, one-way ANOVA). Consequently, *SAMMSON* knock-down was associated with a significant decrease of in cell viability in 92.1 and OMM1 (Supplemental Fig 3B, p<0.0001, one-way ANOVA). Taken together, these results reveal an important role for *SAMMSON* in maintaining UM and CM cell survival *in vitro*.

### SAMMSON inhibition affects protein synthesis and mitochondrial function in UM

To investigate the mechanism by which *SAMMSON* contributes to UM tumor cell survival, we studied *SAMMSON* interaction partners in UM cell lines. First, we evaluated *SAMMSON* binding to p32 and XRN2, 2 interaction partners previously identified in skin melanoma(8,9), using RIP-qPCR. Immunoprecipitation of both p32 and XRN2 in 92.1 cells revealed a 4- and 64-fold enrichment of *SAMMSON* RNA, respectively, indicating that *SAMMSON* binding to these factors is conserved in uveal melanoma cells (Supplemental Fig 4B). We then applied chromatin isolation by RNA purification and mass spectrometry (ChIRP-MS) in 2 UM cell lines (OMM1 and OMM2.3) to identify additional interaction partners of *SAMMSON* in UM cells. We verified enrichment of *SAMMSON* RNA upon *SAMMSON* pull down with biotinylated probes (Supplemental Fig 4A) and subsequently applied mass spectrometry to quantify protein abundance in both the *SAMMSON* and lacZ pull-down samples. We detected 83 and 84 proteins that were significantly enriched upon *SAMMSON* pull down in OMM1 and OMM2.3, respectively, of which 57 were found in both cell lines (Supplemental Fig 4C and Supplemental Table 1). We next performed pathway enrichment analysis and found that pathways involved in mitochondrial translation were highly enriched among the candidate interaction partners in both cell lines (Fig 3C and Supplemental Table 1). Closer inspection of the candidate interaction partners revealed 13 (OMM1) and 7 (OMM2.3) mitochondrial ribosomal proteins (MRPs) (Fig 3A) that are all part of the 39S large mitoribosomal subunit involved in mitochondrial translation. Together with validated p32 and XRN2 interactions, these results suggest a role for *SAMMSON* in regulating translation and mitochondrial function in uveal melanoma. To further investigate the effect of *SAMMSON* knockdown on translation, we quantified global translation levels using a puromycin incorporation assay (SUnSET(19)). *SAMMSON* knockdown significantly impaired translation rates by 56% and 61% in UM cell lines OMM1 and 92.1, respectively (Fig 3B and Supplemental Fig 3C, p=0.0155 (OMM1) and p=0.0077 (92.1), two-way ANOVA).

**Fig 3.**
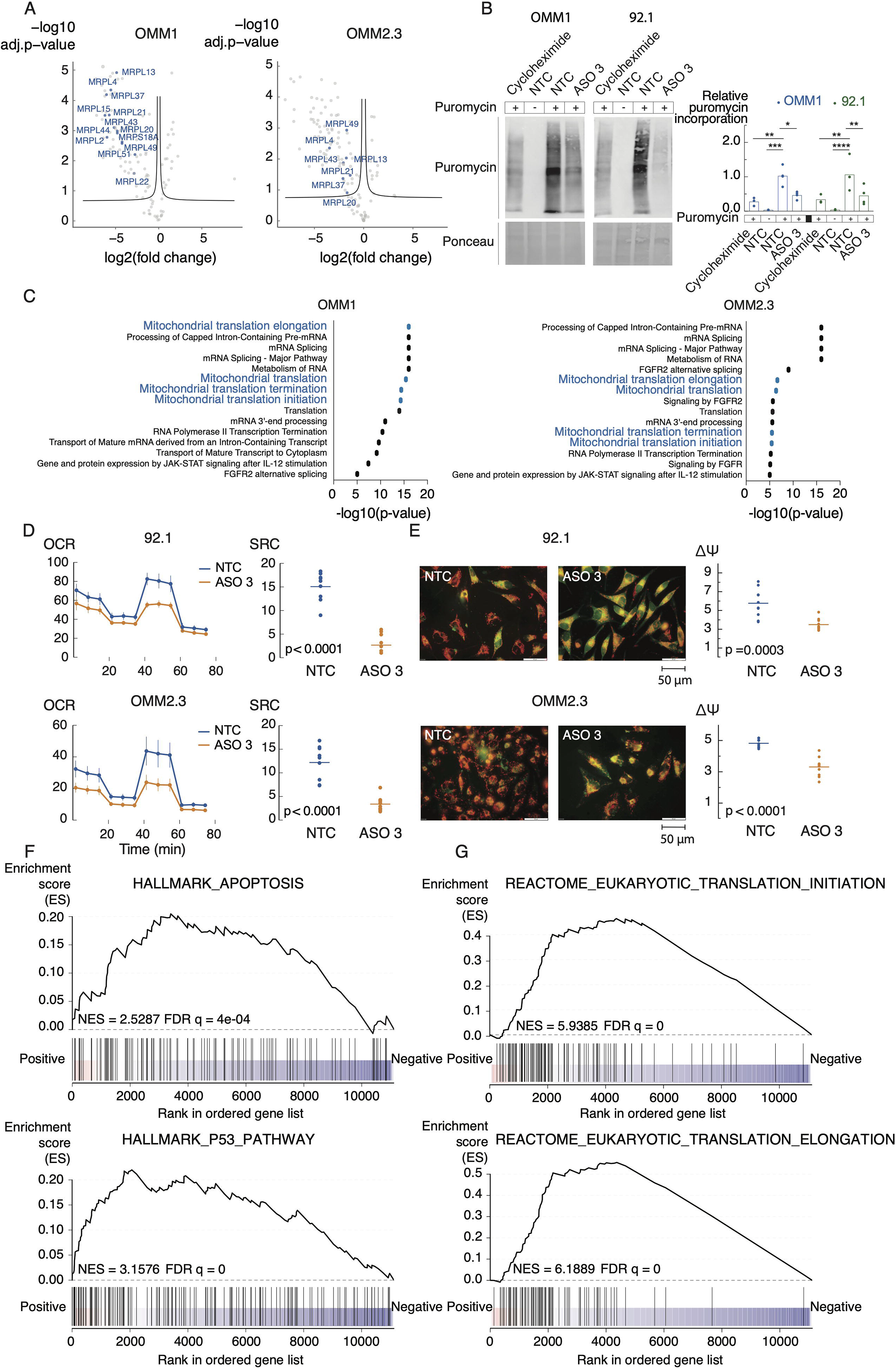
*SAMMSON* inhibition affects global translation and mitochondrial function. **A.** Volcano plot depicting *SAMMSON* interacting proteins identified by ChIRP-MS (left). Probes targeting LacZ were included as a control (right). Mitochondrial ribosomal proteins (MRPs) are indicated on the graph. Significance was calculated using two-sided t-test. **B.** Representative images of WB-SUnSET analysis of UM cells treated with cycloheximide (translation inhibitor, positive control), scrambled ASO (NTC) (without puromycin, negative control), NTC ASO or ASO 3 (50 nM). Quantification of protein synthesis measured by calculating the intensity of the puromycin signal on WB. n=3 replicates. P-values were calculated using two-way ANOVA with Tukey’s multiple comparisons test. **C.** Pathway enrichment analysis for ChIRP results showed participation of *SAMMSON* in mitochondrial translation pathways. **D.** Oxygen Consumption Rate (OCR) measurements over time after sequential injections of oligomycin, fluoro-carbonyl cyanide phenylhydrazone (FCCP) and rotenone/antimycin A in UM cell lines treated for 24 h with NTC ASO or ASO 3 (100 nM). Data are represented as the mean of 3 replicates ± s.d. Spare respiratory capacity (SRC) was obtained by subtracting the basal respiration from the maximal respiration. P-values were calculated using unpaired t-test. **E.** 5,5,6,6-Tetraethylbenzimidazolyl-carbocyanine iodide (JC-1) staining in UM cells treated with with NTC ASO or ASO 3 (100 nM) (magnification of the images x400). Quantification of the electric membrane potential (ΔΨ) as the red over green fluorescence of 10 random selected fields. P-values were calculated using unpaired t-test. **F.** GSEA results for Hallmark gene sets apoptosis and p53 (MsigDB) that are enriched upon ASO 3 treatment (50 nM). Normalized enrichment score (NES) and false discovery rate (FDR) are depicted on the enrichment plot. **G.** GSEA results for c2 curated gene sets translation initiation and translation elongation (MsigDB) which are enriched upon ASO 3 treatment (50 nM). Normalized enrichment score (NES) and false discovery rate (FDR) are depicted on the enrichment plot.

We then explored the importance of *SAMMSON* for proper mitochondrial function by measuring the oxygen consumption rate (OCR) as a proxy for mitochondrial respiration. Differences in OCR after injections of oligomycin, fluoro-carbonyl cyanide phenylhydrazone (FCCP) and rotenone/antimycin A were measured(20). Upon *SAMMSON* knockdown, we observed a significant decrease in mitochondrial spare respiratory capacity (SRC) of 82% and 72% in 92.1 and OMM2.3, respectively (Fig 3D, p<0.0001, unpaired t-test). This was further verified by a 5,5′,6,6′-tetraethylbenzimidazolyl-carbocyanine iodide (JC-1) fluorescence staining, where the electric membrane potential (ΔΨ) is quantified based on the relative red over green fluorescence(17). *SAMMSON* knockdown decreases ΔΨ with 39% and 31% in 92.1 and OMM2.3, respectively, indicating oxidative phosphorylation (OXPHOS) impairment (Fig 3E, p=0.0003 (92.1) and p<0.0001 (OMM2.3), unpaired t-test). These data suggest that the mechanism by which *SAMMSON* affects translation is conserved between UM and SKCM, resulting in an impairment of mitochondrial function. To assess if inhibition of mitochondrial translation using bacteriostatic antibiotics of the tetracyclines family(21–24) can phenocopy *SAMMSON* knock-down, we treated 92.1 and OMM1 UM cells with tigecycline(14). In both cell lines, we observed a tigecycline dose-dependent reduction in cell confluence and induction of apoptosis (Supplemental Fig 5A, p<0.0001 (confluence) and p<0.0001 for all concentrations, except 3.125 μM in 92.1 and p≤0.01 for the two highest concentrations in OMM1 (apoptosis), one-way ANOVA), verifying that UM cells strongly depend on mitochondrial translation. While the growth of non-*SAMMSON* expressing cells HEK293T and CT5.3hTERT is also affected by tigecycline treatment (Supplemental Fig 5B, p<0.0001 (HEK293T) and p≤0.05 for all concentrations, except 3.125 μM (CT5.3hTERT), two-way ANOVA), tigecycline was markedly less effective in CT5.3hTERT cells (cell viability reduction of 35% using 50 μM of tigecycline compared to ≥55% in 92.1, OMM1 and HEK293T).

To assess the impact of *SAMMSON* on the UM transcriptome, we performed shallow RNA sequencing of UM cell lines 92.1 and OMM1 treated with either NTC ASO or ASO 3. Differential gene expression analysis revealed 378 up- and 255 downregulated genes (adjusted p-value <0.05). Gene set enrichment analysis (GSEA) revealed a significant upregulation of gene sets associated with apoptosis and p53 response upon *SAMMSON* silencing (Fig 3F, FDR q=4e-04 (apoptosis) and FDR q=0 (p53) and Supplemental Table 2), supporting our phenotypic observations. Surprisingly, *SAMMSON* inhibition also resulted in a transcriptional activation of genes involved in translation, suggesting a negative feedback loop in response to the inhibition of translation (Fig 3G, FDR q=0).

### SAMMSON inhibition suppresses UM PDX growth

*In vivo* anti-tumor effects of *SAMMSON* knockdown were evaluated in three independent experiments using two UM patient derived xenografts (PDX) models (MP46 (GNAQ^Q209L^) and MEL077 (derived from a patient progressing on the immune checkpoint inhibitor pembrolizumab)) showing high *SAMMSON* expression levels (Fig 1C). When the tumor xenografts reached a volume of 60-180 mm^3^, the mice were randomly separated into two groups and were subcutaneously injected with either NTC ASO or ASO 3 (10 mg/kg). Tumor growth was significantly delayed (Fig 4B, p=0.0001 at endpoint, two-way ANOVA) and tumor weight significantly reduced in MEL077 PDX mice treated with ASO 3 compared to NTC ASO (Fig 4C-D, p=0.0183, unpaired t-test). These observations were validated in an independent experiment using the same PDX MEL077 model (Supplemental Fig 6C) as well as in UM PDX model MP46 (Fig 4A, p=0.0050 at endpoint, two-way ANOVA). No weight loss of the mice was observed after treatment, suggesting that the ASO 3 dose and administration regimen did not result in major toxic side effects (Supplemental Fig 6A, B, D).

**Fig 4.**
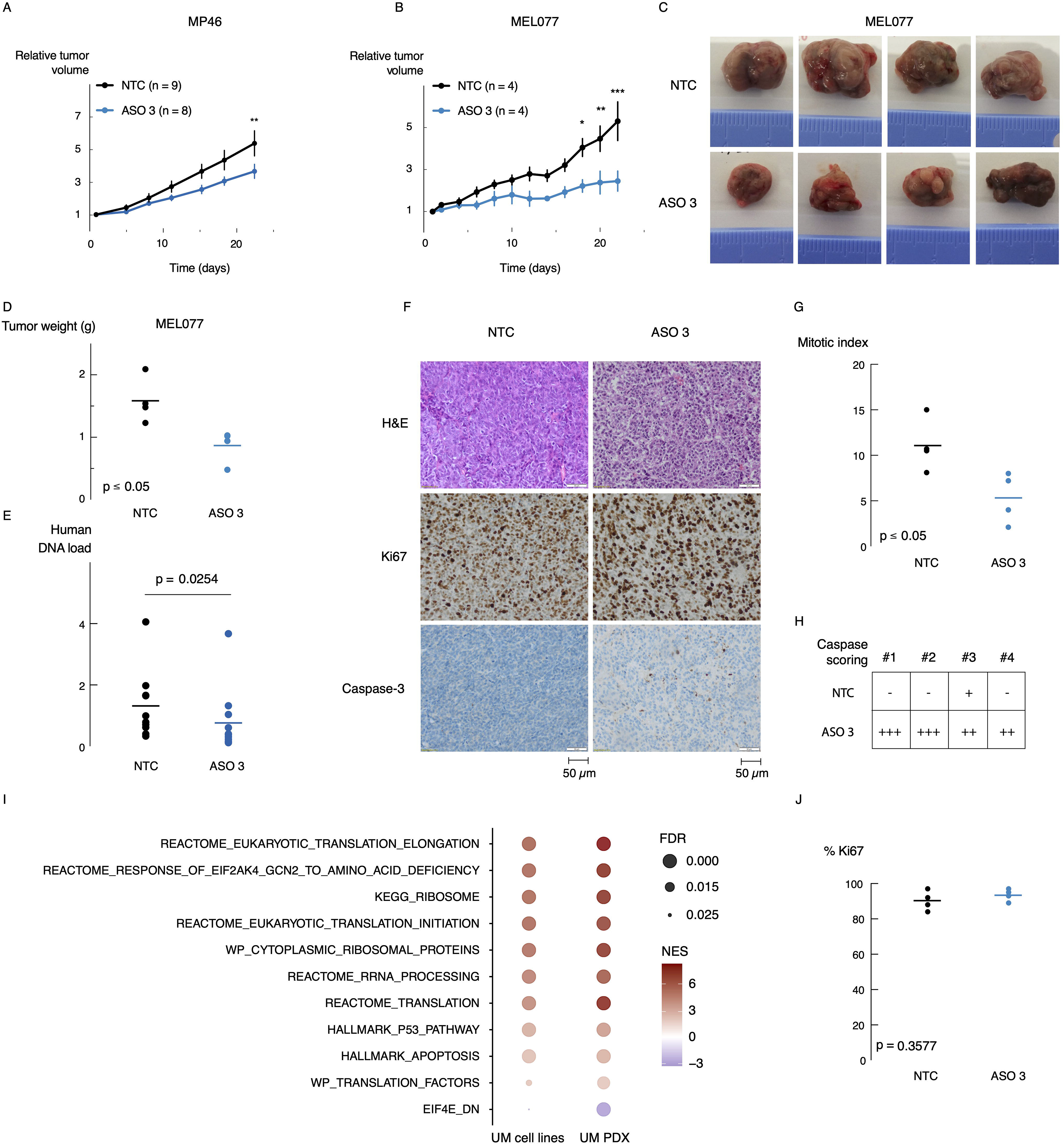
*SAMMSON* inhibition reduces tumor growth *in vivo*. **A.** Relative tumor volume of MP46 PDX mice subcutaneously injected with scrambled ASO (NTC) or ASO 3 (10 mg/kg). Data are mean ± s.e.m. of multiple replicates (n=8-9). P-values were calculated using two-way ANOVA (** p≤0.01). **B.** Relative tumor volume of MEL077 PDX mice subcutaneously injected with NTC ASO or ASO 3 (10 mg/kg). Data are mean ± s.e.m. of multiple replicates (n=4). P-values were calculated using two-way ANOVA (* p≤0.05, ** p≤0.01, *** p≤0.001). **C.** Representative tumors of MEL077 PDX mice 22 days after treatment. **D.** Tumor weight of tumors as shown in **C.** P-value was calculated using unpaired t-test. **E.** Human DNA load measured in lung tissues of MEL077 (n=4) and MP46 (n=10-11) mice by means of qPCR of Alu-Sq repetitive sequence. P-value was calculated using unpaired t-test. **F.** H&E, Ki67 and Caspase-3 staining of MEL077 tumor sections 22 days after treatment with NTC ASO or ASO 3. **G.** Number of mitotic cells based on H&E staining in F of 10 random high power fields (HPFs) (magnification x400). Data are represented as the mean of 10 HPFs per tumor sample (n=4). P-value was calculated using unpaired t-test. **H.** Caspase-3 scoring based on morphology and staining intensity as shown in F of 4 tumor samples treated with either NTC ASO or ASO 3. **I.** Selected GSEA results of RNA seq data from UM cell lines OMM1 and 92.1 and UM MEL077 PDX tumors demonstrating overlapping enrichment of gene sets involved in apoptosis, p53 and translation upon ASO 3 treatment. The depth of the color represents the normalized enrichment score (NES). The area of the circle represents the false discovery rate (FDR). **J.** Percentage of Ki67 positive cells in 500 counted cells per tumor sample (n=4) (magnification x100). P-value was calculated using unpaired t-test.

Immunohistochemistry (IHC) analysis with anti-Ki67 of the MEL077 model revealed highly aggressive tumors, with over 80% of proliferating cells in all tumor samples (Fig 4F). While the difference in Ki67 expression between the ASO 3 and NTC ASO treated mice was too small to be statistically significant (Fig 4F, J, p=0.3577, unpaired t-test), further investigation of the mitotic index showed a significant reduction in the number of mitotic cells in the ASO 3 treated tumor samples (Fig 4F, G, p=0.0278, unpaired t-test). In addition, caspase-3 activity was elevated in tumors from mice treated with ASO 3 compared to NTC ASO, indicating *SAMMSON* knock-down induces apoptosis *in vivo* (Fig 4F, H). Moreover, all tumor samples were correctly labeled as ‘ASO 3-treated’ or ‘NTC ASO-treated’, independently by two pathologists and blinded to treatment.

To further investigate these tumors at the molecular level, we applied RNA-sequencing to generate transcriptome profiles to define pathways significantly enriched or repressed upon PDX ASO treatment. Notably, we observed a strong overlap (p<0.0001, Fisher exact test) between pathways induced in the PDX model and pathways induced in the UM cell lines upon *SAMMSON* knockdown. Gene sets related to apoptosis and p53 response were significantly upregulated (Fig 4I, FDR q<0.05, Supplemental Table 2). Similarly, pathways related to translation were also found transcriptionally upregulated.

Although no macro-metastatic lesions were observed in both PDX models, we investigated the presence of tumoral DNA in blood and various murine tissues resected from both PDX models(25). To this end, we extracted genomic DNA (gDNA) from blood, liver and lung of PDX mice that received ASO 3 or NTC ASO treatment and performed quantitative PCR using the human specific Alu-Sq, SVA and LINE-1 repetitive sequences as an indicator of tumor load in these tissues. Copy number levels of the murine Hprt1 and Pthlh genes were used for normalization. Tumor DNA was only detected in the murine lung tissue samples and compared to NTC ASO treated mice, mice treated with ASO 3 had significantly reduced levels of tumor DNA in lung tissue (Fig 4E and Supplemental Fig 6E, F, p=0.0254 (Alu-Sq), p=0.0357 (LINE-1), p=0.0202 (SVA), unpaired t-test).

## Discussion

Long non-coding RNAs are crucial players in many cellular and biological processes including regulation of gene expression, cell growth, differentiation and development. Aberrant expression of lncRNAs is observed in virtually all tumor types and some lncRNAs appear to be cell type- and tissue-specific(26,27). The lncRNA *SAMMSON* was discovered as a melanoma specific lncRNA(8), but several studies have also shown occasional expression in gastric cancer (GC), hepatocellular carcinoma (HCC), glioblastoma (GBM) and papillary thyroid carcinoma (PTC)(28–32). In the present study, we found that *SAMMSON* is expressed in more than 80% of uveal melanoma tumors at levels that are substantially higher compared to non-melanoma tumors. In addition, *SAMMSON* expression could be observed in conjunctival melanoma (CM) cells that are genetically and phenotypically more related to skin melanoma. While 50% of uveal melanoma tumors are characterized by loss of an entire copy of chromosome 3, *SAMMSON* expression levels are not reduced in monosomy 3 tumors whereas the majority of genes on chromosome 3 do show a clear gene-dosage effect. These observations suggest the existence of a compensation mechanism that requires further investigation. Potentially, monosomy 3 tumors upregulate one or multiple transcription factors that drive *SAMMSON* expression from the remaining allele.

The lack of treatments for UM patients with metastatic dissemination that occurs in 50% of the patients and is accompanied by extremely poor survival(4) demonstrates the high unmet need for new treatment modalities. Therapeutic nucleic acid-based approaches like siRNA and antisense oligonucleotides (ASO) hold enormous potential to target RNA molecules, including lncRNAs. Advances in therapeutic ASO and siRNA technology research, such as the development of locked nucleid acid (LNA) and S-constrained ethyl (cEt) modified ASOs and 2’-OMe and 2’-fluoro modified siRNAs, enabled the development and approval of several ASO and siRNA drugs to treat diseases such as spinal muscular atrophy (SMA) and hereditary transthyretin-mediated amyloidosis (hATTR) (33–35). Our work demonstrates that, similar to skin melanoma, ASO-mediated *SAMMSON* knockdown in various *in vitro* and *in vivo* UM models induces a potent anti-tumor response including growth reduction, induction of apoptosis, and reduced levels of tumor-derived DNA in lung tissue. Whether this DNA is cell-free or derived from tumor cells invading these tissues remains to be investigated. The fact that we did not observe tumor DNA in blood may be indicative of the latter. Additional studies in uveal melanoma xenograft models that metastasize (36–38) should further assess the impact of *SAMMSON* knockdown on metastatic disease. Our *in vitro* data, based on multiple cell lines with a different genetic background, and *in vivo* data, based on two genetically different UM PDX models, suggest that the genetic background of the tumor does not influence the response to *SAMMSON* inhibition, further highlighting the broad relevance of *SAMMSON* inhibition as a potential therapeutic strategy.

Earlier studies in SKCM demonstrated *SAMMSON* interaction with proteins involved in mitochondrial and cytosolic translation, such as p32, XRN2 and CARF(9). In UM, interactions with p32 and XRN2 were confirmed and novel candidate interactions with multiple proteins belonging to the 39S large mitoribosomal subunit were identified, further supporting the role of *SAMMSON* in regulating mitochondrial and cytosolic translation. Additional mechanistic studies are required to further elucidate the role of these novel candidate interactors. Of note, XRN2 and p32 were not identified by mass-spectrometry analysis, suggesting that we may have missed other interaction partners. In support of these interactions and the established role of *SAMMSON* in SKCM, *SAMMSON* inhibition effectively impairs global translation rates and consequently mitochondrial translation, since all proteins required for mitochondrial translation, including the mitoribosomal proteins and the mitochondrial translation factors, are translated by cytosolic ribosomes(39). All 13 polypeptides synthesized by the mitochondrial ribosomes (mitoribosomes) are essential components of the oxidative phosphorylation machinery(40). *SAMMSON* inhibition indeed impairs mitochondrial function, as shown in two UM cell lines. This results in mitochondrial precursor overaccumulation stress (mPOS), characterized by the toxic accumulation and aggregation of unimported mitochondrial proteins in the cytosol(41). Cells need to balance between import of mitochondrial proteins and the cytosolic capacity to handle unimported mitochondrial proteins. Induction of mPOS can tip that balance, hereby compromising cell viability. Although *SAMMSON* inhibition affects global translation, transcriptome analysis in both UM cell lines and PDX models revealed a transcriptional upregulation of various components involved in the translational machinery. These observations suggest a negative feedback response to global translation inhibition, which has also been observed when treating cells with translation inhibitors such as cycloheximide (CHX) (42).

In conclusion, our work shows that Inhibition of *SAMMSON* impairs translation, which consequently affects mitochondrial function and ultimately results in decreased cell viability and induction of apoptosis, both *in vitro* and *in vivo*. Together, our findings suggest that *SAMMSON* is an attractive therapeutic target for UM and that *SAMMSON* can be targeted *in vivo* using ASO technology.

## Acknowledgements

We would like to thank the VIB proteomics core for performing LC-MS/MS analysis and Prof. Olivier De Wever for providing the CT5.3hTERT cell line.

The authors would also like to thank the Animal PlatformCRP2-UMS 3612 CNRS-US25 Inserm-IRD (Faculty of Pharmacy, Paris University, Paris), and the Animal Platform of Institut Curie. They also thank Justine Fleury for her technical assistance (Laboratory of preclinical Investigation, Institut Curie).

---

**Supplemental Fig 1.**
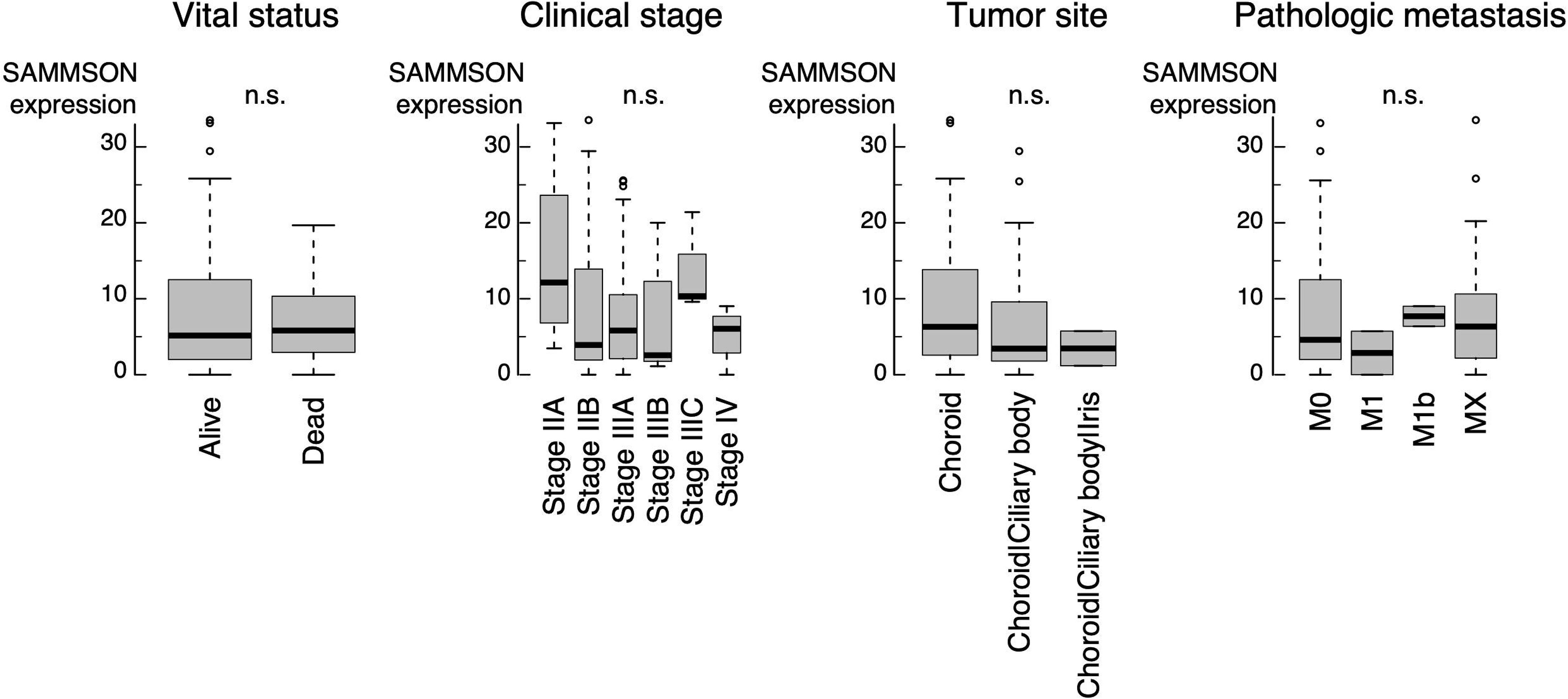
*SAMMSON* expression is independent from patient survival, tumor stage, tumor localization site and metastatic state. RNA sequencing data from >10 000 tumor samples and 32 cancer types (TCGA) showing no correlation between *SAMMSON* expression and the vital status, clinical stage, tumor site or pathological metastatic state of the UM patient. n.s. = not significant.

**Supplemental Fig 2.**
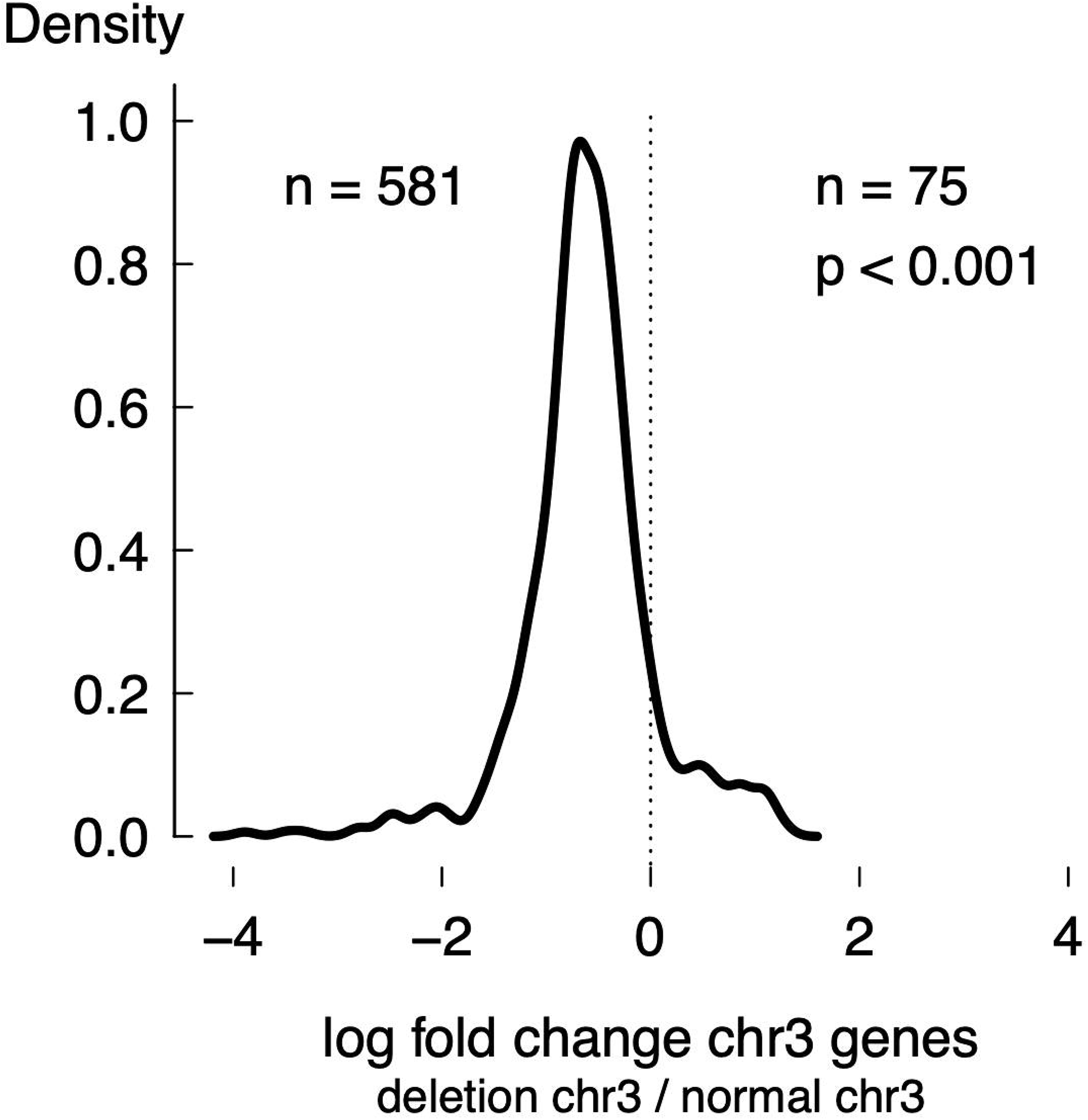
Expressed genes in monosomy 3 UM tumors compared to disomy 3 UM tumors. 656 genes are expressed in monosomy 3 tumors of which 581 (89%) are downregulated and 75 (11%) upregulated. P-value was calculated using Mann-Whitney test.

**Supplemental Fig 3.**
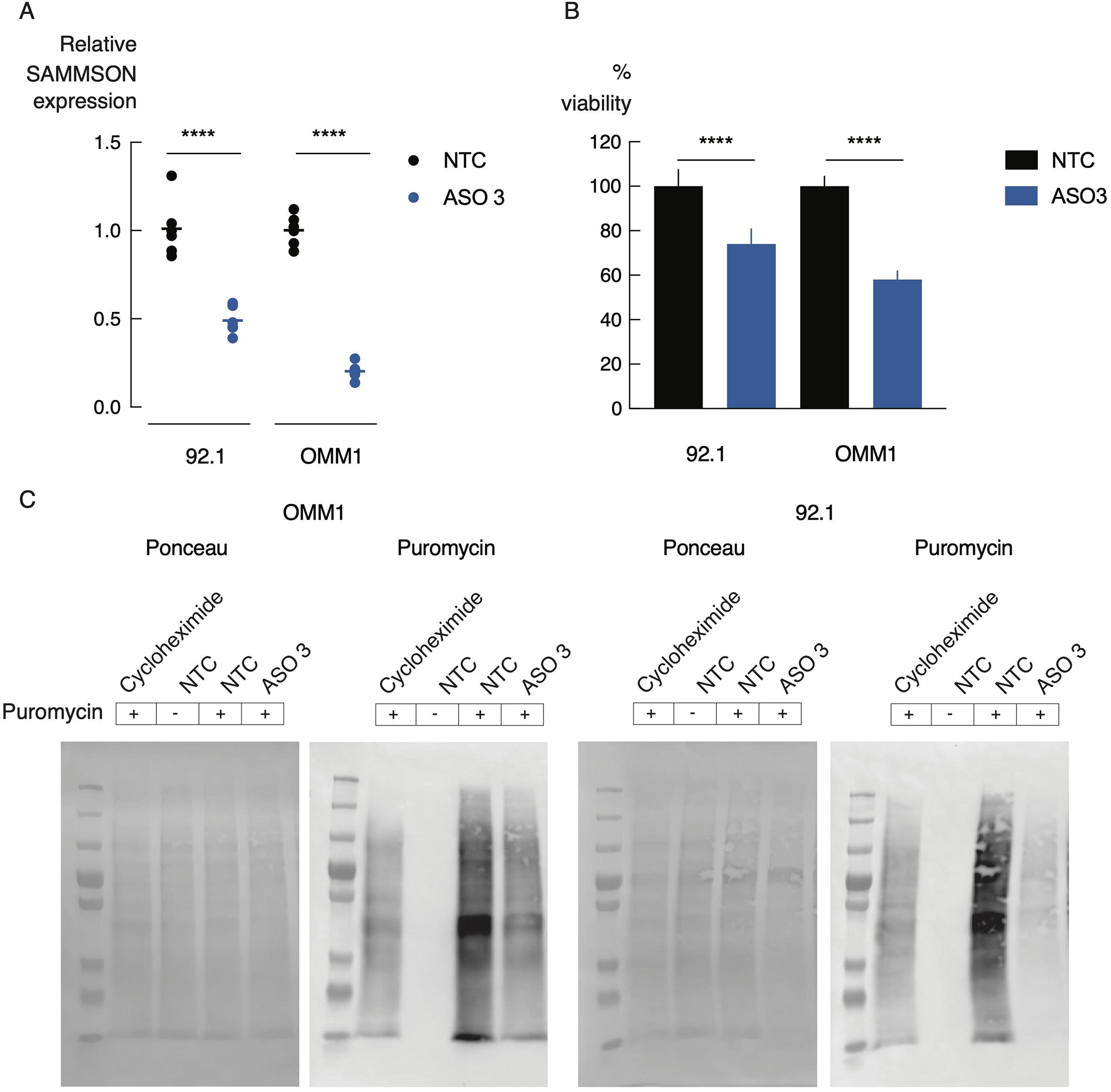
Non-lipid-based delivery of ASOs in UM cells. **A.** Relative *SAMMSON* expression in two UM cell lines transfected with scrambled ASO (NTC) or ASO 3 (100 nM) using the TransIT-X2 transfection reagent. The results are obtained 48 h after transfection and represent the average of six replicates. P-values were calculated using one-way ANOVA with Tukey’s multiple comparisons test (**** p≤0.0001) B Viability (Cell titer glo) assay in NTC ASO and ASO 3 treated UM cells using the TransIT-X2 transfection reagent (72 h post-transfection). The data are presented as the mean of 10 replicates ± s.d. P-values were calculated using one-way ANOVA with Tukey’s multiple comparisons test (**** p≤0.0001). **C.** Uncropped images of WB-SUnSET analysis of UM cells treated with cycloheximide (translation inhibitor, positive control), scrambled ASO (NTC) (without puromycin, negative control), NTC ASO or ASO 3 (50 nM).

**Supplemental Fig 4.**
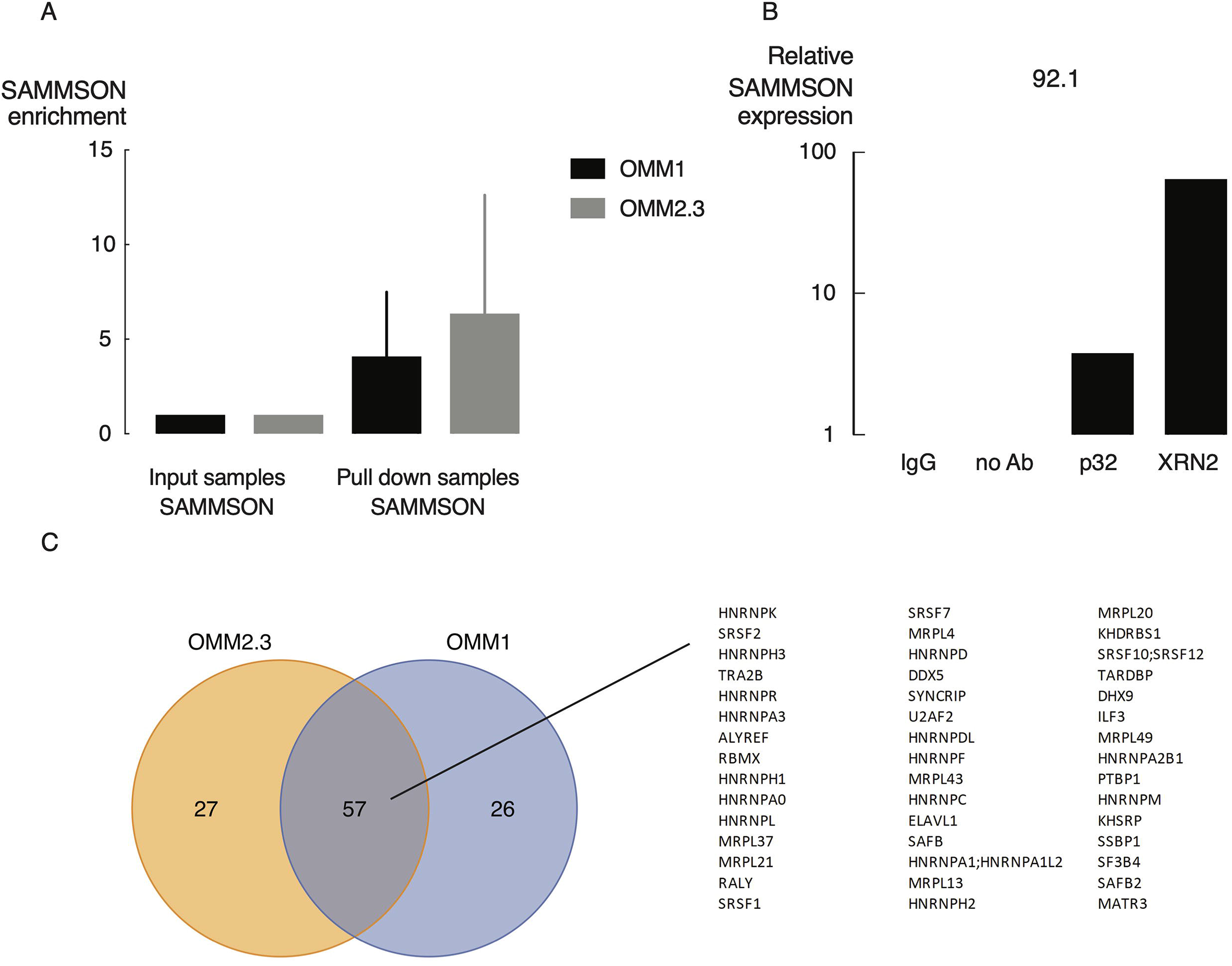
Identification of *SAMMSON* interaction partners. **A.** Enrichment of *SAMMSON* RNA upon *SAMMSON* pull down with biotinylated probes in ChIRP-MS. **B.** p32 and XRN2 were identified as *SAMMSON* interacting proteins by means of RIP-qPCR. **C.** Significantly enriched proteins upon *SAMMSON* pull down in two UM cell lines with 57 overlapping proteins.

**Supplemental Fig 5.**
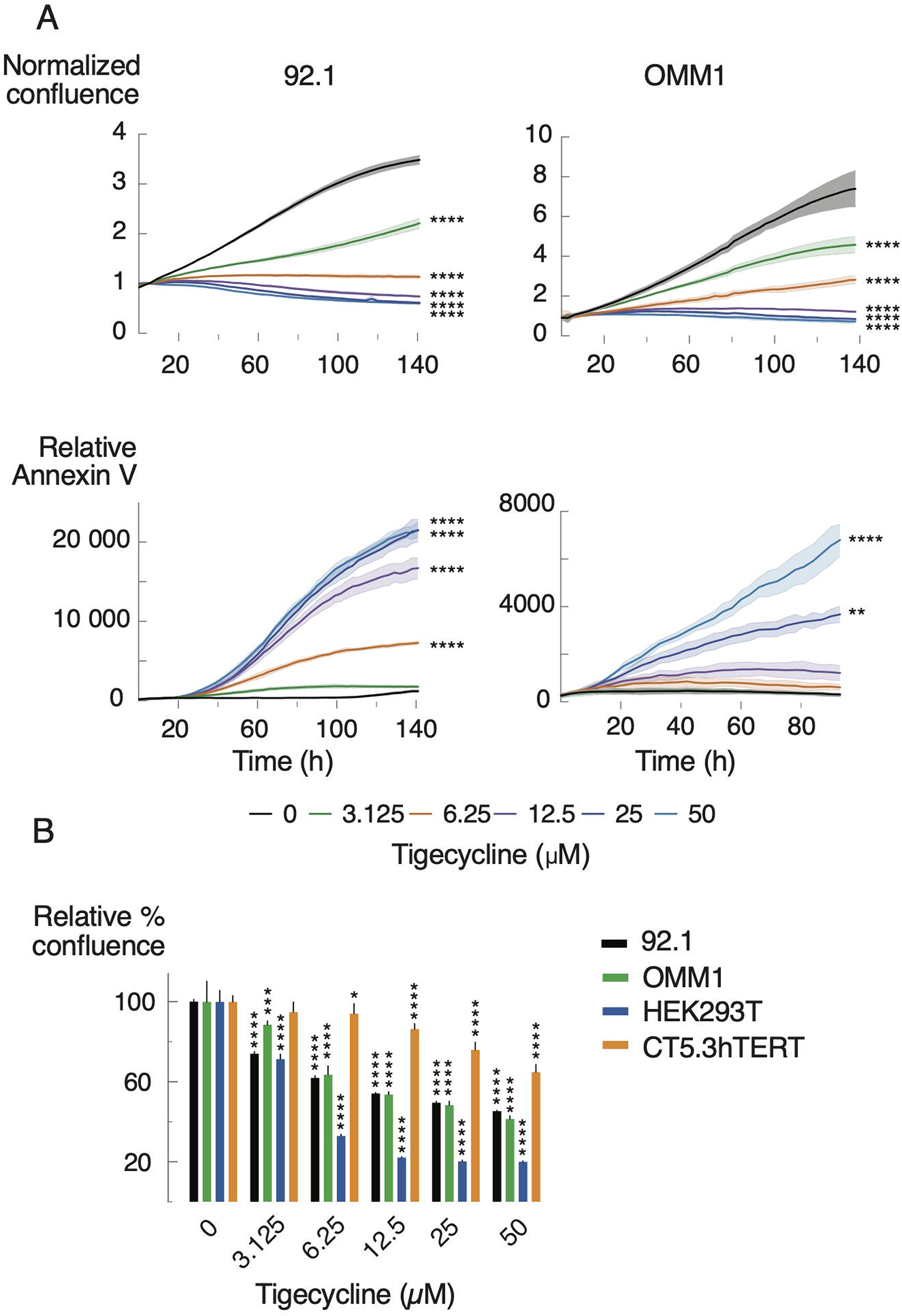
Tigecycline phenocopies *SAMMSON* inhibition in UM cells. **A.** Reduction of proliferation (confluence) and induction of apoptosis (annexin V) in UM cell lines 92.1 and OMM1 upon treatment with multiple concentrations of tigecycline (3.125, 6.25, 12.5, 25 or 50 μM) compared to untreated cells (0 μM) measured with time-lapse microscopy (every 2-3 h) using the IncuCyte device. **B.** Relative confluence 48 h after treatment with multiple concentrations of tigecycline (3.125, 6.25, 12.5, 25 or 50 μM) compared to untreated cells (0 μM). Error bars represent ± s.d. of 5 replicates. P-values are calculated using one-way ANOVA with Dunnett’s multiple comparisons test (last time point for A), * p≤0.05, ** p≤0.01, *** p≤0.001, **** p≤0.0001).

**Supplemental Fig 6.**
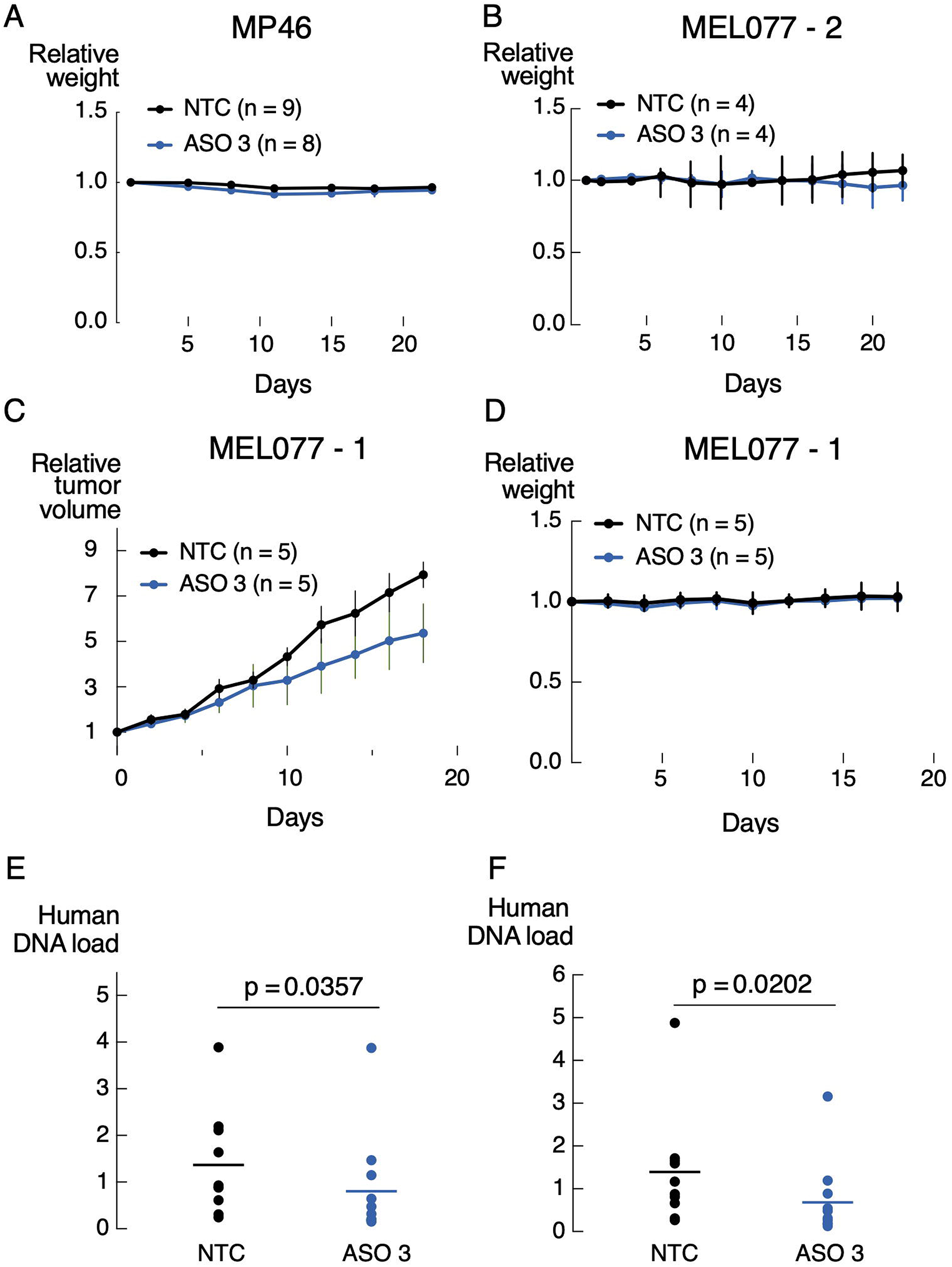
*SAMMSON* inhibition slows down tumor growth *in vivo*. **A, B, D.** Relative tumor weight of MP46 PDX mice (A) or MEL-077 PDX mice (B, D) treated with scrambled ASO (NTC) or ASO 3 (10 mg/kg). Data are mean ± s.e.m. of multiple replicates (n=8-9 (MP46), n=4 (MEL077–2, n=5 (MEL077–1)) **C.** Relative tumor volume of MEL-077 PDX mice subcutaneously injected with NTC ASO or ASO 3 (10 mg/kg). Data are mean ± s.e.m. of multiple replicates (n=5). **E, F.** Human DNA load measured in lung tissues of MEL077 (n=4) and MP46 (n=10-11) mice by means of qPCR for LINE-1 (E) or SVA (F) repetitive sequence. P-value was calculated using unpaired t-test.

